# Engineering ligand stabilized aquaporin reporters for magnetic resonance imaging

**DOI:** 10.1101/2023.06.02.543364

**Authors:** Jason Yun, Logan Baldini, Yimeng Huang, Eugene Li, Honghao Li, Asish N. Chacko, Austin D.C. Miller, Jinyang Wan, Arnab Mukherjee

## Abstract

Imaging transgene expression in live tissues requires reporters that are detectable with deeply penetrant modalities, such as magnetic resonance imaging (MRI). Here, we show that LSAqp1, a water channel engineered from aquaporin-1, can be used to create background-free, drug-gated, and multiplex images of gene expression using MRI. LSAqp1 is a fusion protein composed of aquaporin-1 and a degradation tag that is sensitive to a cell-permeable ligand, which allows for dynamic small molecule modulation of MRI signals. LSAqp1 improves specificity for imaging gene expression by allowing reporter signals to be conditionally activated and distinguished from the tissue background by difference imaging. In addition, by engineering destabilized aquaporin-1 variants with different ligand requirements, it is possible to image distinct cell types simultaneously. Finally, we expressed LSAqp1 in a tumor model and showed successful in vivo imaging of gene expression without background activity. LSAqp1 provides a conceptually unique approach to accurately measure gene expression in living organisms by combining the physics of water diffusion and biotechnology tools to control protein stability.

## INTRODUCTION

Genetic reporters, commonly based on fluorescent and bioluminescent proteins, allow gene expression to be monitored with outstanding specificity in living systems. However, optical imaging in vertebrates is challenging because of absorption and scattering of light by thick tissues. In contrast to optical methods, magnetic resonance imaging (MRI) provides unfettered depth access and enables wide-field imaging with fairly high spatial resolution. To make MRI signals fully genetically encodable, we recently introduced metal-free reporters based on the human water channel aquaporin 1 (Aqp1)^1^. Aqp1 expression increases the rate of water exchange across the cell membrane, which permits its detection using an MRI technique known as diffusion-weighted imaging. This technique applies a weighting factor (*b*) determined by magnetic field gradients, such that the MRI signal intensity (*S*) scales with the diffusivity of water molecules (*D*) as *S* ∝ *e* (−*b* · *D*)^2,3^. Cells expressing Aqp1 allow for faster diffusion of water, making them appear hypointense (darker) compared to background tissue. Recent studies have applied Aqp1 in conjunction with diffusion-weighted MRI to trace neural connectivity^4^, observe tumor gene activity^5^, and image astrocytes on a brain-wide scale in live mice^6^.

Unlike conventional reporter gene readouts, biological water diffusion is sensitive to elements of tissue structure such as cell size, extracellular volume fraction, tissue geometry, and subcellular architecture^7,8^. Therefore, the specificity of using Aqp1 as a molecular reporter is potentially decreased in milieu where changes in tissue structure, for example, due to cell death or swelling, could alter water diffusivity in ways that compete with or cancel the diffusion enhancement induced by Aqp1. Furthermore, air-tissue interfaces (e.g., near the amygdala in mice) and calcified tissue also lead to signal darkening in MRI, which may be difficult to distinguish from Aqp1-derived contrast. To ensure specific, accurate, and reliable imaging with Aqp1, we need an approach to resolve Aqp1 signals from the tissue background and other nonspecific factors that alter water diffusion.

One approach for distinguishing reporter signals from the tissue background in MRI is differential imaging. This approach involves subtracting images acquired with the reporter in an “on” state from those obtained when the reporter is “off” to create a background-free hotspot that reflects signals specific to reporter gene expression. For example, reporters based on chemical exchange saturation transfer (CEST) are visualized as difference signals obtained by saturating the probe at a unique resonance frequency (offset from that of bulk water) and subtracting pre and post-saturation images^9–13^. MRI reporters based on gas vesicles also permit background-free imaging by subtracting images acquired with intact vesicles from images obtained after acoustically collapsing the vesicles, which destroys their contrast^14^. Although the MRI signals from CEST reporters and gas vesicles can be dynamically modulated for background-free imaging, these reporter classes have important limitations. CEST provides limited sensitivity, while gas vesicles involve large multi-gene clusters that are challenging to express in mammalian cells. Conversely, Aqp1 is a single-gene, human-origin reporter that is a highly sensitive but lacks a built-in modulation mechanism to differentiate its signals from nonspecific effects.

An ideal approach for modulating Aqp1 signals should achieve ample on-off amplitude, function in multiple cell types and in vivo, exhibit a favorable safety profile, and avoid off-target effects, particularly on endogenously expressed aquaporins. To this end, we sought to chemically modulate Aqp1 stability by incorporating ligand-dependent degrons. Degrons are protein motifs, that when fused to a protein of interest, recruit ubiquitin ligase, resulting in ubiquitination and degradation of the tagged protein^15,16^. Ligand-dependent degrons rely on small molecules to modulate protein degradation^17^.

Ligand binding can either stabilize a target protein by blocking degron-ubiquitin ligase interactions^18–21^ or can facilitate the recruitment of ubiquitin ligase, leading to the target protein’s degradation^22–27^. Both methods permit dynamic regulation of protein function, and are widely used as genetic tools in basic research, drug discovery^28^, and synthetic biology^29–33^. Here, we exploit ligand-dependent degrons to image gene expression without background interference, through small-molecule induced changes in Aqp1 stability. We engineered and characterized six degron-tagged Aqp1 reporters and identified an optimal construct (LSAqp1) for small-molecule control of Aqp1 signals. We demonstrate that LSAqp1 permits chemically resolved background-free imaging in multiple cell types and subcutaneous tumors in mice. We further show that degron-tagging of Aqp1 can be exploited to unlock two additional capabilities of considerable interest for reporter gene applications – notably, drug-gated monitoring of transcriptional activity and imaging distinct cell types in multiplex.

## RESULTS

### Degron-tagging of Aqp1 enables background-free imaging by small-molecule modulation

To determine whether degron-tagging permits small-molecule modulation of Aqp1, we tested six ligand-degron pairs, three each for triggering ligand-induced degradation (LID) and ligand-induced stabilization (LIS) of the protein target^18–20,23–25,27^. We cloned each degron at the C-terminus of Aqp1 and used lentiviral transduction to stably express the resulting constructs in Chinese hamster ovary cells (CHO). We treated the engineered cells with their respective degron-matched ligands and used diffusion-weighted MRI to quantify the percent change in the diffusion coefficients of ligand-treated cells relative to the vehicle-treated controls (𝛥*D*/*D*_*o*_) (**Supplementary Fig. 1**). The LIS fusions generated a larger fold-change in diffusivity compared to the LID tags (**Fig. 1a-b**), with the largest response measured in cells expressing Aqp1 fused to a degron derived from a F36V/L106P double mutant of the mammalian prolyl isomerase, FKBP12^18,34^. Hereafter, we refer to the Aqp1-FKBP12^F36V/L106P^ construct as LSAqp1 (Ligand-Stabilized Aqp1) for brevity. Treatment of cells expressing LSAqp1 with the cognate ligand shield-1 elicited an 86 ± 6 % (mean ± s.d., *n* = 9, *P* < 10^−9^, 2-sided t-test) increase in the diffusion coefficient relative to vehicle-treated cells (**Fig. 1b**), with half-maximum activation occurring in approximately five hours (**Fig. 1c**). Shield-1 was non-toxic (**Fig. 1d**) and specific to LSAqp1 with no effect on diffusion signals in wild-type cells, erythrocytes (that natively express aquaporin-4), and cells engineered to express wild-type Aqp1 without a degron tag (**Fig. 1e, Supplementary Fig. 2**). We further examined the ability of shield-1 to modulate LSAqp1 in distinct cell-types, including Jurkat (T cells), U87 (glioblastoma), and RAW264.7 (macrophage). As before, we used lentiviral transduction to stably express LSAqp1 in each cell line and measured shield-1 dependent changes in the diffusion coefficients. We observed a significant response in all cell types, with ΔD/D_o_ ranging from 56 ± 3 % in RAW264.7 to 189 ± 21 % in Jurkat cells (*n* = 4, *P* < 0.002, 2-sided t-test) (**Fig. 1f**).

**Figure 1:**
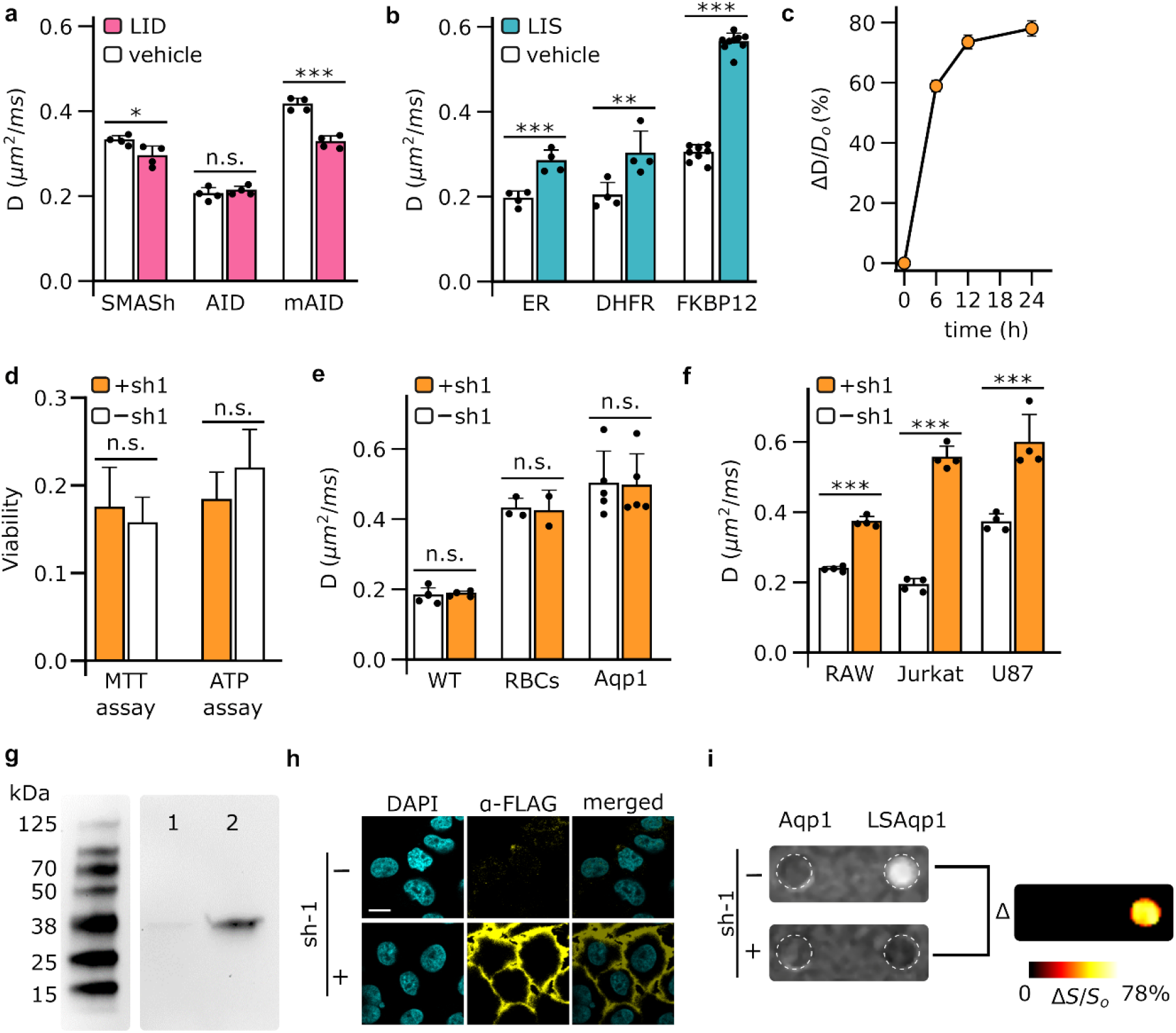
Engineering small-molecule modulated Aqp1 for background-free imaging. **a**, Fusing ligand-induced degradation (LID) tags to the C-terminus of Aqp1 generates modest changes in the diffusion coefficients of ligand-treated CHO cells relative to vehicle-treated controls. **b**, In contrast, fusing ligand-induced stabilization (LIS) tags to the C-terminus generates substantial changes in the diffusivities of ligand-treated CHO cells relative to controls. The largest response was observed in cells expressing Aqp1 fused to the FKBP12F36V/L106P degron (hereafter, the chimeric construct is referred to as LSAqp1), which was stabilized by a small-molecule, shield-1 (denoted as sh-1 in figures). **c**, Time-dependent increase in diffusion rate of LSAqp1-transduced cells following treatment with shield-1 (1 µM). **d**, Treatment of LSAqp1-transduced cells with shield-1 did not result in acute toxicity, as determined by MTT and ATP-based viability assays. **e**, Shield-1 modulation is specific to LSAqp1 and does not affect the baseline diffusivity of wild type cells or the enhanced diffusivity of red blood cells or cells engineered to express aquaporin-1. **f**, LSAqp1 permits small-molecule modulation of diffusivity in multiple cell types. **g**, Immunoblotting with anti-FLAG antibodies reveals a band of the expected size of LSAqp1 in membrane fractions extracted from shield-1 treated CHO cells (lane 2), which was only faintly observed in extracts obtained from vehicle-treated cells (lane 1). **h**, Confocal microscopy using anti-FLAG antibodies revealed membrane-localized LSAqp1 in shield-1 treated, but not vehicle-treated CHO cells. Images of both cells under both treatment conditions were acquired using identical optical settings (see Methods) and were displayed on the same color-scale. Scale bar is 5 µm. **i**, Background-free MRI with LSAqp1. Differential imaging was performed by voxel-wise subtraction of diffusion-weighted images acquired with and without shield-1 treatment. The ensuing image was denoised using a median filter and displayed as a pseudo-colored “hotspot”. Error bars represent the standard deviation (*n* ≥ 3 biological replicates). * denotes *P* < 0.05, ** denotes *P* < 0.01, *** denotes *P* < 0.005, and n.s. is non-significant (*P* ≥ 0.05). *P*-values were computed based on 2-sided Student’s t-test (unpaired).

To verify that shield-1 modulation of diffusion coefficients correlates with stabilization of LSAqp1, we inserted a FLAG epitope at the N-terminus and used an anti-FLAG antibody to immunoblot membrane fractions prepared from shield-1 and vehicle-treated cells. In agreement with the MRI-based diffusion measurements, we observed a prominent band in extracts prepared from shield-1 treated cells, which was absent in the control cells (**Fig. 1g, Supplementary Fig. 3**). We further performed immunofluorescence imaging to probe shield-1 dependent changes in membrane-localized LSAqp1. As the N-terminus of LSAqp1 is located in the cytoplasm, we inserted a FLAG epitope in an extracellular loop between Gln43 and Thr44, which allowed us to probe LSAqp1 without having to permeabilize the cells. Consistent with the Western blotting data, immunofluorescence imaging also revealed membrane-localized LSAqp1 largely in shield-1 treated cell lines (**Fig. 1h**).

To probe the mechanism by which the FKBP12^F36V/L106P^ degron destabilizes Aqp1 in the absence of shield-1 we tested whether degradation could be rescued by treating cells with inhibitors of each of the two major protein degradation pathways: chloroquine, an inhibitor of lysosomal degradation, and MG132, an inhibitor of proteasomal degradation. We used MRI to compare diffusion coefficients of untreated LSAqp1-transduced cells to those of cells treated with either inhibitor. We observed a partial recovery of diffusivity (ΔD/D_o_ = 25 ± 5 %, *n* = 7, *P* = 0.01, 2-sided t-test) in the chloroquine-treated cohort, suggesting that LSAqp1 is partly degraded by the lysosomal pathway (**Supplementary Fig. 4**), although other mechanisms of LSAqp1 modulation, such as membrane trafficking, cannot be ruled out. Since degradation is influenced by the terminus at which the degron is attached, we further wondered whether moving the FKBP12^F36V/L106P^ degron to the N-terminus of Aqp1 would confer a similar shield-1 dependent change in diffusivity. We found the performance of N-terminal fusion to be cell-type dependent, exhibiting a larger fold-change than the C-terminal fusion in CHO cells, a smaller response in U87 and Jurkat cells, while being non-responsive in RAW264.7 cells (**Supplementary Fig. 5**). We focused the rest of our study on the C-terminal fusion (viz., LSAqp1), which exhibited a more consistent performance across different cell types.

Based on the favorable characteristics of LSAqp1, we hypothesized that this reporter would permit background-free imaging of engineered cells via small-molecule modulation of MRI signal. To test the concept, we acquired diffusion-weighted images of LSAqp1-transduced CHO cells alongside cells expressing degron-free Aqp1 and red blood cells (which express Aqp4). We housed the cells in an agarose phantom to mimic background tissue and performed voxel-wise subtraction of diffusion-weighted images acquired with and without shield-1 incubation. Through this approach, we were able to clearly distinguish the LSAqp1 cells from both the agarose background and other cellular signals (**Fig. 1i, Supplementary Fig. 6**). This approach can also be used to image LSAqp1 cells by calculating difference signals from diffusion maps, using each voxel’s diffusion coefficient instead of intensity (**Supplementary Fig. 7**).

### LSAqp1 enables gated imaging of transcriptional activity

Having established small-molecule modulation of LSAqp1, we wondered whether this approach could be applied for small molecule-gated imaging of transcriptional activity. Reporters whose output is gated by drug exposure enable the monitoring of gene expression specifically during user-defined epochs, such as a treatment regime or behavioral task^35–37^. The design of such reporters exploits logical AND operation to generate an output only when promoter activity is accompanied by an external trigger applied by the experimenter^38,39^ (**Fig. 2a, Supplementary Fig. 8**). To test whether LSAqp1 permits chemically gated imaging of transcriptional activity, we designed a construct in which LSAqp1 expression was driven by a doxycycline inducible promoter (**Fig. 2b**). For comparison, we also constructed a similar vector using the conventional (ungated) Aqp1. We delivered each construct to CHO cells by lentiviral infection, treated stable cells with both shield-1 and doxycycline, as well as each compound individually, and acquired whole-cell diffusion measurements as before. In contrast to Aqp1, where promoter activity (induced by doxycycline) alone is sufficient to generate a reporter gene output (**Fig. 2c**), both promoter induction and shield-1 are needed to elicit maximum response in LSAqp1 (ΔD/D_o_ = 95 ± 19 %, *P* = 1.8 × 10^−4^, *n* = 6, 2-sided t-test) (**Fig. 2d**). In agreement with these results, differential imaging revealed an LSAqp1 signal only when promoter activity was accompanied by shield-1 application (**Fig. 2e, Supplementary Fig. 8**). Taken together, these results indicate the potential of using LSAqp1 as a small-molecule gated MRI reporter to capture transcriptional activity in a user-selected window defined by exposure to shield-1.

**Figure 2:**
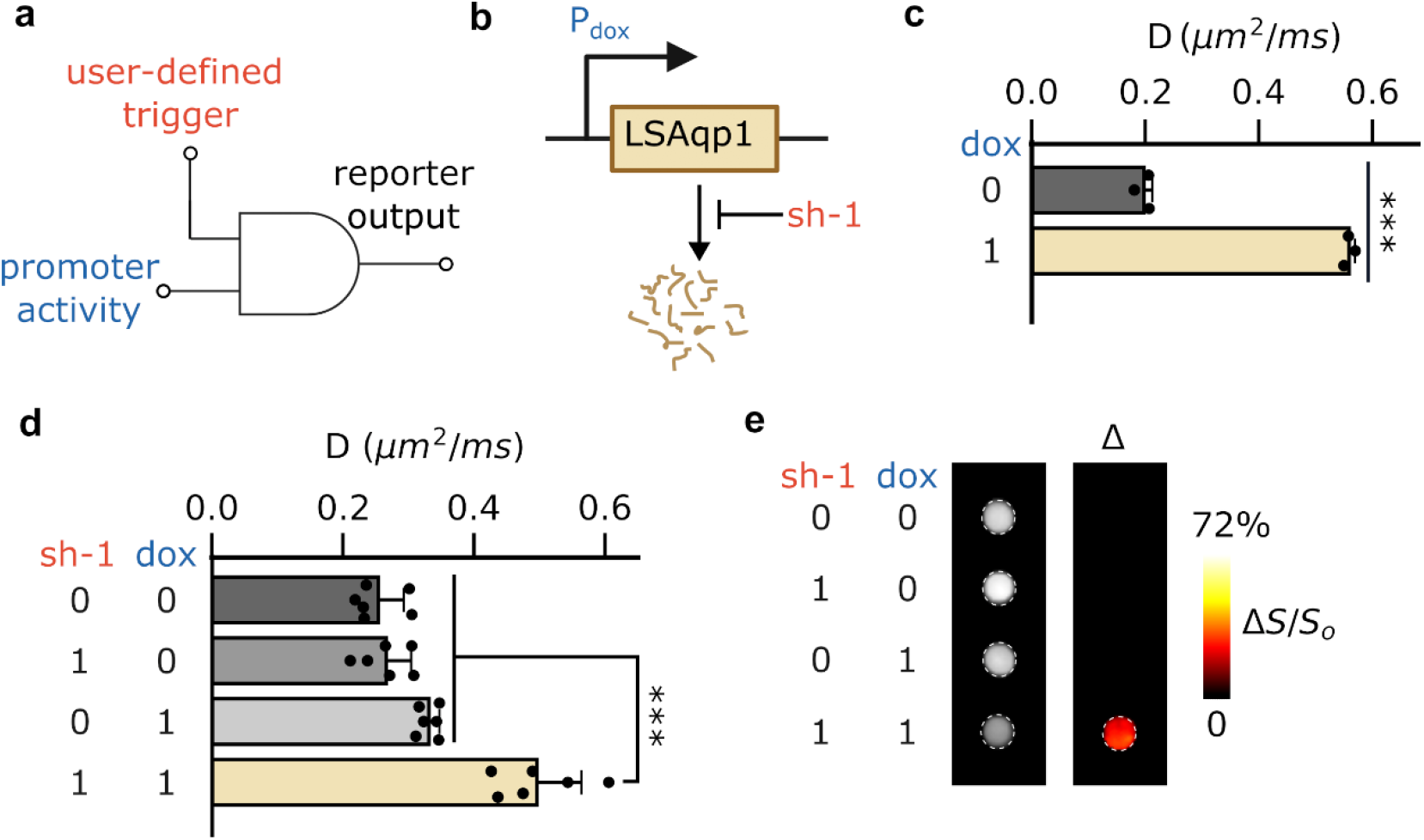
Drug-gated imaging of promoter activity. **a**, Chemically gated reporters leverage genetic AND circuits to generate an output only when transcriptional activity is accompanied by an external trigger. **b**, Reporter construct for drug-gated imaging of promoter activity induced by doxycycline. **c**, In conventional reporters such as Aqp1, promoter activity alone is sufficient to drive the reporter gene. **d**, In the gated reporter based on LSAqp1, both promoter activity and shield-1 were required to elicit the maximum response. **e**, A positive differential signal was observed when transcriptional activity (induced by doxycycline) is accompanied by shield-1. Differential imaging was performed by subtracting the diffusion-weighted images of untreated CHO cells (sh-1 = 0, dox = 0) from the images of cells treated with either shield-1 (sh-1 = 1, dox = 0), doxycycline (sh-1 = 0, dox = 1), or both chemicals simultaneously (sh-1 = 1, dox = 1). The ensuing signal was denoised by median filtering and displayed in pseudo-color. Error bars represent the standard deviation (*n* = 6 biological replicates). *** denotes *P* < 0.005. *P*-values were computed based on 2-sided Student’s t-test (unpaired).

### Degron-tagged reporters allow substrate resolved imaging of cells in multiplex

Many applications in biomedicine would benefit from the ability to use MRI-based reporter genes to simultaneously map multiple biological features, such as distinct cell types or promoter activities. Although proton MRI is fundamentally a “monochromatic” technique, we hypothesized that it would be possible to use LSAqp1 in conjunction with a second degron-tagged Aqp1 reporter to image cells in multiplex by stabilizing one reporter at a time and performing differential imaging. For this approach to work, degron-tagged Aqp1 reporters should exhibit non-overlapping ligand requirements, thereby permitting stepwise modulation of diffusivity with ligand addition. Conceptually, this approach is inspired by substrate-resolved imaging in bioluminescence^40–42^ and CEST^13^, where reporter signals are separated based on their selectivity for orthogonal ligands. We first established that the three ligand-stabilized Aqp1 constructs identified in our initial screening, namely LSAqp1, Aq-dhfr (dihydrofolate reductase degron), and Aq-ER (human estrogen receptor-based degron) are mutually orthogonal, that is, cells transduced with either of the three reporters do not respond to non-cognate ligands (**Fig. 3a, Supplementary Fig. 8**). To demonstrate “2-color” imaging, we prepared a 1:1 mixture of CHO cells transduced with LSAqp1 and Aq-dhfr. We obtained whole-cell diffusion measurements on untreated cells, cells treated with shield-1 only, and cells treated with both shield-1 and trimethoprim. We observed an expected stepwise increase in the diffusion coefficient with the addition of ligands (**Fig. 3b**). Next, we performed pairwise subtraction of diffusion-weighted images obtained after treating cells with one or both ligands, which allowed the two reporter-expressing cell types to be distinguished in the mixed population (**Fig. 3c**). To verify that the above approach is generalizable to other cell types and degron-tagged constructs, we also demonstrated multiplex imaging of a U87 + Jurkat cell mixture engineered to respectively express LSAqp1 and Aq-ER (**Fig. 3d,e**).

**Figure 3:**
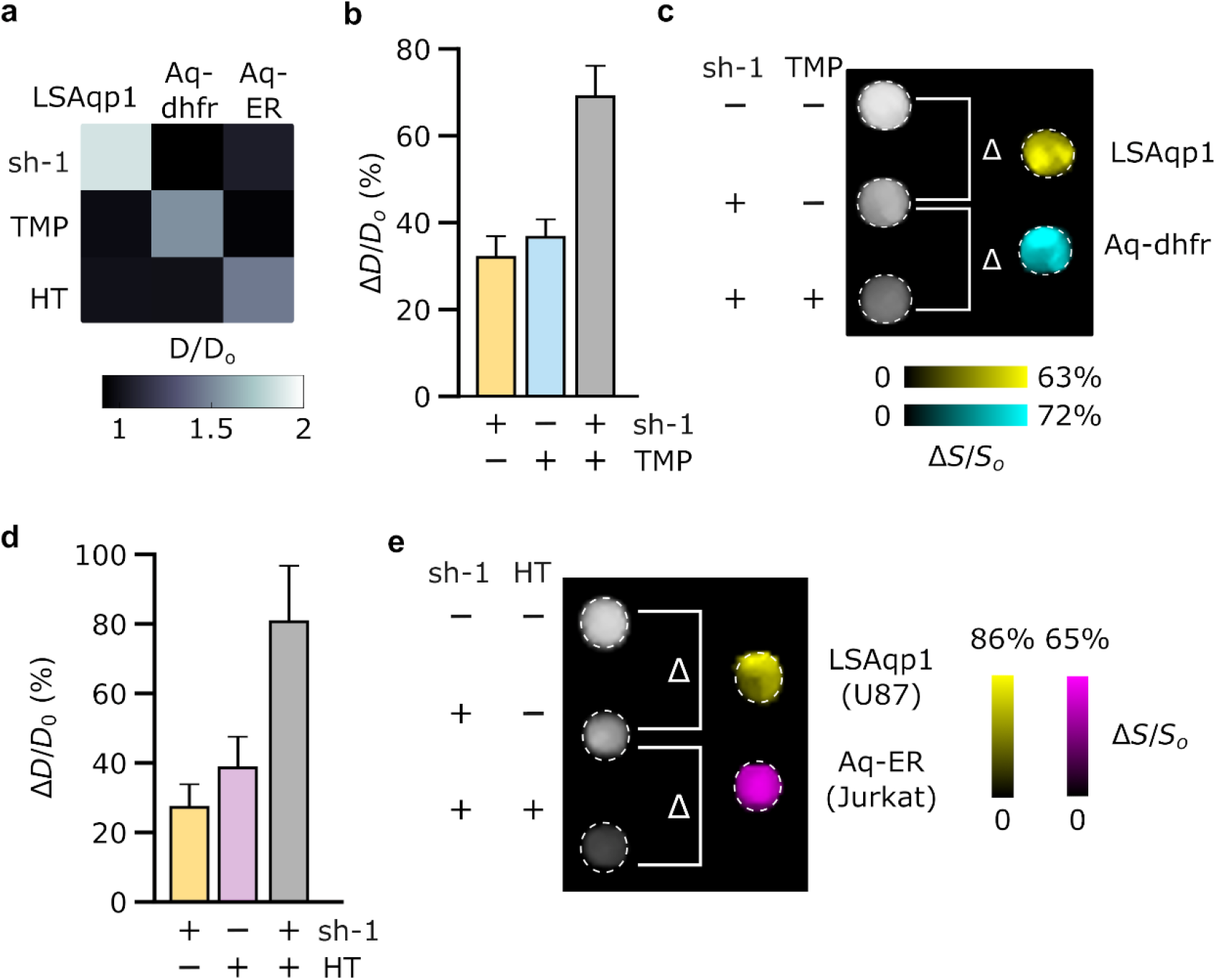
Unmixing of Aqp1-based diffusion signals for multiplex imaging. **a**, LSAqp1 (responsive to shield-1, sh-1), Aq-dhfr (stabilized by trimethoprim, TMP), and Aq-ER (responsive to 4-hydroxytamoxifen, HT) are mutually orthogonal, because treatment with non-cognate ligands does not increase the diffusion coefficients of CHO cells transduced with the respective reporter construct. **b**, Ligand-dependent increase in diffusion (relative to vehicle-treated cells) in a mixed population comprising equal numbers of LSAqp1- and Aq-dhfr transduced cells. Ligand addition permits the stepwise modulation of diffusivity by stabilizing one or both degron-tagged Aqp1 constructs. **c**, Components of the mixed-cell population were resolved by differential imaging. Subtraction of diffusion-weighted images of untreated cells from images acquired after treatment with shield-1 reveals the LSAqp1 population. Likewise, subtracting diffusion-weighted images of shield-1 treated cells from those obtained after treatment with shield-1 and TMP revealed the Aq-dhfr population. The images were denoised by median filtering and pseudo-colored to distinguish the LSAqp1- and Aq-dhfr-expressing sub-populations. **d**, Ligand-dependent increase in the diffusion coefficient of a mixed population comprising U87 and Jurkat cells transduced respectively with LSAqp1 and Aq-ER. As before, ligand addition permits the stepwise modulation of diffusion. **e**, The two cell types can be distinguished by differential imaging after sequential treatment with ligands. The images were denoised by median filtering and pseudo-colored to distinguish between the two cell-types. Error bars represent standard deviation (*n* ≥ 4 biological replicates).

### LSAqp1 enables background-free imaging in vivo

Finally, to evaluate the in vivo utility of LSAqp1, we used the reporter in a mouse tumor xenograft model, which provides a common testing ground for new reporter gene techniques. We formed transgenic tumors by injecting CHO cells bilaterally into the hind flanks of mice: one set of cells was engineered to express LSAqp1 and the other Aqp1-only, viz., without an N-terminal degron (**Fig. 4a**). After the tumors reached a volume of 100-300 mm^3^, we acquired two sets of diffusion-weighted images: the first before injecting shield-1 and the next six hours post-injection. In agreement with the trends observed in vitro (**Fig. 1b,f**), the LSAqp1 tumors showed a statistically significant increase in the diffusion coefficient following shield-1 injection (ΔD/D_o_ = 62.5 ± 20 %, *n* = 5 mice, *P* = 0.004, 2-sided t-test), while the diffusivity of the Aqp1 tumors was not altered (*n* = 5 mice, *P* = 0.41, 2-sided t-test) (**Fig. 4b**). Accordingly, small-molecule modulation allows LSAqp1 expression to be specifically visualized by differential imaging and overlaying the ensuing signal on a higher-resolution anatomical image (**Fig. 4c)**. A similar approach can also be applied to generate differential images based on diffusion maps, where each voxel is represented by the computed diffusion coefficient (**Fig. 4d**).

**Fig. 4.**
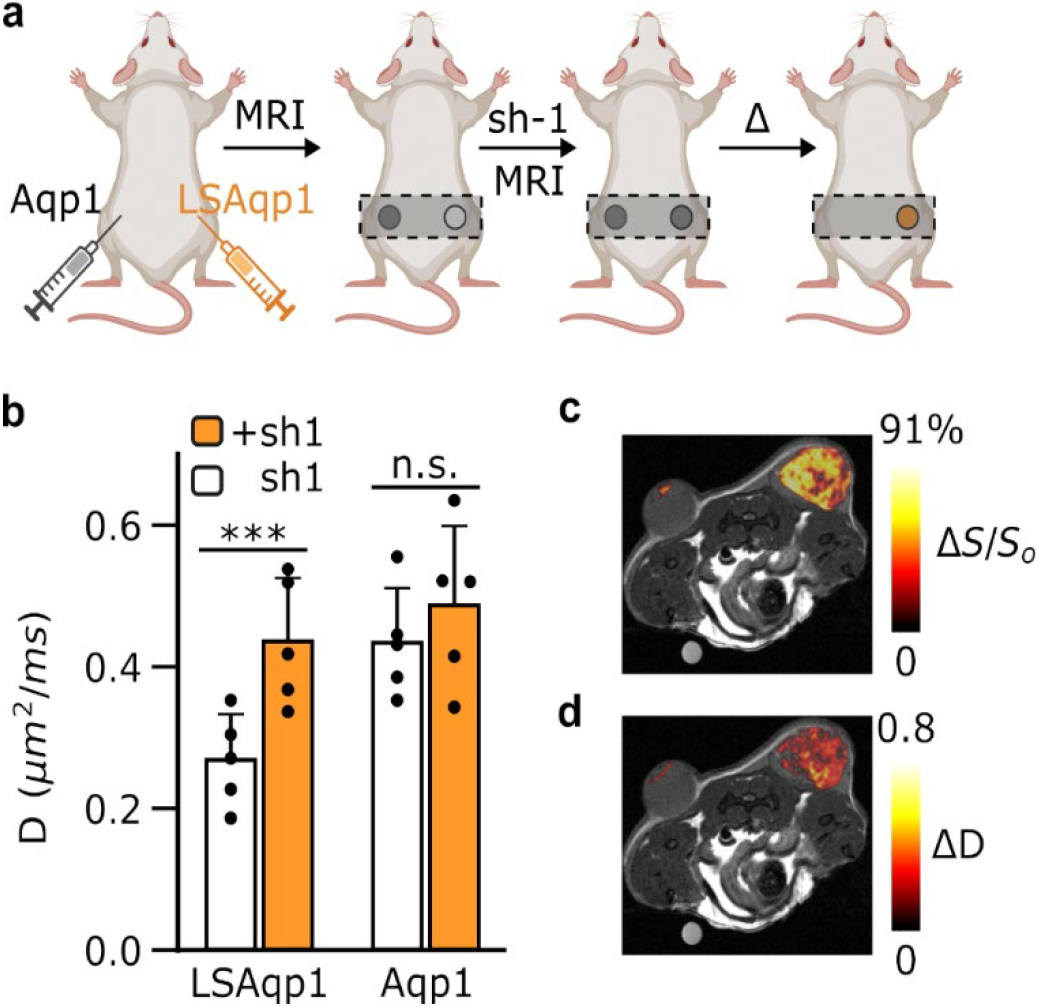
In vivo tumor imaging using LSAqp1. **a**, Schematic outline of the in vivo study. **b**, Diffusion coefficients in transgenic tumors engineered to express LSAqp1 or Aqp1 (*n* = 5) measured before and after intraperitoneal injection of shield-1 (10 mg/kg). A post-injection increase in diffusion was observed only in the LSAqp1-expressing tumor cells. **c**, Small-molecule modulation with shield-1 allows the LSAqp1 tumor to be specifically visualized by subtracting post-injection diffusion-weighted images from pre-injection images and overlaying the difference on an anatomical image. A water-filled phantom was placed near the mouse to monitor the accuracy and consistency of MRI measurements. **d**, Differential imaging can also be implemented based on diffusion maps where each voxel is represented by its measured diffusion rate. Error bars represent the standard deviation. *** denotes *P* < 0.005, and n.s. is non-significant (*P* ≥ 0.05). *P*-values were computed based on 2-sided Student’s t-test (paired).

## DISCUSSION

Our work establishes ligand-induced stabilization of Aqp1 as a conceptually new approach for imaging gene expression using MRI. The best-performing construct, LSAqp1, enables small-molecule modulation of diffusion signals using a safe, cell-permeable, and bioorthogonal chemical, shield-1. Differential imaging erases signals unrelated to the reporter, thereby making LSAqp1 expression visible as a background-free “hotspot”. LSAqp1 also permits the detection of promoter activity via genetic AND circuits, paving the way for time-locked monitoring of transcriptional signals in user-defined experimental windows. Finally, we showed that orthogonal degron-tagged Aqp1 pairs may be paired for multiplex imaging of gene expression by chemical unmixing of diffusion-weighted signals.

We anticipate that LSAqp1 will be broadly useful for applications requiring gene expression to be accurately tracked in non-transparent animals. Notably, the in vivo safety and biodistribution of shield-1 is already established in various rodent models^43–45^ and furthermore, shield-1 was recently shown to cross the blood-brain barrier^46^. Accordingly, LSAqp1 can be used to monitor viable cells and map gene expression in tumor tissue, unambiguously separating reporter signals from factors like necrosis, apoptosis, and edema, which cause nonspecific diffusion increases in the tumor microenvironment. LSAqp1 also offers the advantage of tracking transient changes in transcriptional activity, which would be missed by conventional reporters with long half-lives^47^. Additionally, the ability to gate LSAqp1 output will be useful for neuroscience applications to accurately correlate brain-wide gene expression patterns with behavior and neuromodulation paradigms. Recent studies showing that first-generation Aqp1 functions as a viable reporter in the mouse brain^4,6^ provide an encouraging indication for extending LSAqp1 for large-scale neural mapping studies. Other potential applications of LSAqp1 include tracking the location of immune- and stem cell-based therapies inside the body. Finally, with continued improvements in sensitivity, degron-tagged Aqp1(s) can be used in multiplex to monitor multiple cell types and transcriptional signals in living organisms.

We envision at least three avenues for improving LSAqp1. First, although the kinetics of LSAqp1 modulation aligns with many biological processes, such as tumor growth, cell migration, and gene expression, faster modulation would allow access to a broader range of biological functions and simplify the imaging workflow by allowing pre- and post-injection images to be acquired in quick succession without moving the animal between scans. Second, increasing the ligand-controlled change in diffusion rate (𝛥*D*/*D*_*o*_) would increase LSAqp1’s sensitivity, allowing for improved detection of rare cell types or weakly expressed genes. We believe that both of these goals can be accomplished through molecular engineering of LSAqp1 using techniques such as directed evolution, which has been successfully applied to optimize ligand-gated degrons in the context of cytosolic proteins. Although the low throughput of MRI poses a practical challenge for directed evolution, our ability to detect shield-1 modulation by epitope tagging of LSAqp1 offers a convenient strategy for screening mutant libraries using high-throughput methods, such as fluorescence- or magnetic-activated cell sorting. Third, while tumor xenografts are widely used as testing grounds for noninvasive reporters, future studies are needed to evaluate the performance of LSAqp1 in additional organs and organ systems.

In summary, we expect that LSAqp1 will synergize with technological advances in diffusion-weighted MRI^48^, the advent of complementary reporter gene modalities^49,50^, and existing functional MRI paradigms such as metabolic imaging^51^, to bring us closer to the goal of exploring complex biological functions on large spatial scales and with molecular and cell-type specificity in intact vertebrates.

## METHODS

### Reagents

Reagents for PCR amplification and Gibson assembly were purchased from New England Biolabs (Ipswich, MA, USA). Polyethyleneimine (linear, 25 kDa) was purchased from Polysciences (Warrington, PA, USA). Lenti-X concentrator was purchased from Takara Bio (Kusatsu, Shiga, Japan). Polybrene was purchased from Santa Cruz Biotechnology (Dallas, TX, US). Dulbecco’s Modified Eagle Media (DMEM), sodium pyruvate, doxycycline hyclate, penicillin-streptomycin (10^4^ units/mL penicillin and 10 mg/mL streptomycin), beta-mercaptoethanol, MG-132, chloroquine diphosphate, polyethylene glycol 400, Tween®-80, and *N,N*-dimethylformamide were purchased from Sigma-Aldrich (St. Louis, MO, USA). Roswell Park Memorial Institute media (RPMI 1640), Gibco™ fetal bovine serum (FBS), Pierce™ BCA Protein Assay Kit, and DAPI (4′,6-diamidino-2-phenylindole) lactate salt were purchased from Thermo Fisher Scientific (Waltham, MA, USA). MycoAlert® Plus Mycoplasma Detection Assay was purchased from Lonza. Trimethoprim and 4-hydroxytamoxifen were purchased from Fisher Scientific; shield-1 was obtained from Aobious (Gloucester, MA, USA); indole-3-acetic acid from Neta Scientific (Hainesport, NJ, USA), asunaprevir was purchased from AdooQ Biosciences (Irvine, CA, USA); and dTag-13 from Tocris Bioscience (Bristol, UK). USP-grade doxycycline hyclate was purchased from Fresenius Kabi USA. Reagents for denaturing polyacrylamide gel electrophoresis, including pre-cast gels, Laemli buffer, Tris-buffered saline with 0.1 % Tween®-20 (TBS-T), non-fat dried milk, and Clarity™ Western ECL substrate were purchased from Bio-Rad (Hercules, CA, USA). Ladders for Western blotting were purchased from LI-COR Biosciences (Lincoln, NE, USA) - WesternSure Chemiluminescent pre-stained ladder or GoldBio (St. Louis, MO, USA) - BLUEstain™ Protein ladder (11-245 kDa). ProteoExtract® native membrane protein extraction kit and 10 kDa Amicon® Ultra-15 centrifugal filters were purchased from Millipore Sigma (USA). Assays for measuring cell toxicity were purchased from Promega (Madison, WI, USA). Anti-FLAG primary antibody was purchased from Sigma-Aldrich (#F1804). Horseradish peroxidase-conjugated (#TL280988) and Alexa Fluor 647 (#A21235) conjugated goat anti-mouse IgG secondary antibodies were purchased from Thermo Fisher. Type F immersion oil was purchased from Leica Microsystems (Deerfield, IL, USA).

### Molecular biology

Plasmids harboring the various degron sequences - DHFR (Addgene 29326), ER (Addgene 37261), miniIAA7 (Addgene 129721), SMASh (Addgene 68853), and FKBP12 (Addgene 17416) were amplified using Q5 High-Fidelity 2X Master Mix and cloned by Gibson assembly in a lentiviral transfer vector at the C or N-terminus of the aquaporin-1 reporter (Aqp1) under the control of either a constitutive promoter, EF1α (Addgene 60058) or a doxycycline-inducible minimal CMV promoter (Addgene 26431). An EGFP reporter was co-expressed with Aqp1 using an internal ribosome entry site (IRES) to allow selection of stably transduced cell by fluorescence-activated cell sorting (FACS). All constructs were verified by Sanger DNA sequencing (Genewiz, San Diego, CA, USA) or whole-plasmid nanopore sequencing (Plasmidsaurus, Eugene, OR, USA).

### Cell culture and engineering

Cells were routinely cultured at 37 °C in a humidified 5 % CO_2_ incubator using DMEM (CHO-TetON, U87, RAW264.7) or RPMI (Jurkat) supplemented with 10 % FBS, 100 U/mL penicillin, and 100 µg/mL streptomycin. For cells harboring an inducible reporter gene cassette, doxycycline hyclate (1 µg/mL) was added to the growth media at the time of seeding. Cells were periodically checked for Mycoplasma contamination using a bioluminescent assay (MycoAlert® Plus). Lentivirus was packaged using a three plasmid system, comprising a packaging plasmid, an envelope plasmid encoding the vesicular stomatitis virus G protein (to pseudo type lentivirus for broad tropism), and the transfer plasmids constructed above. The three plasmids (22 µg each of the packaging and transfer plasmid; 4.5 µg of envelope plasmid) were delivered to 293T cells by transient transfection using polyethyleneimine. Approximately 17-24 h post transfection, the 293T cells were treated with sodium butyrate (10 mM) to promote expression of viral genes. Viral production was allowed to proceed for another 72 h before precipitating the cell-free supernatant using a commercial polyethylene glycol formulation (Lenti-X concentrator) to harvest lentivirus. The resulting viral particles were resuspended in 200 µL DMEM and immediately used to transduce the recipient cell lines, including CHO Tet ON, Jurkat, U87, and RAW 264.7. For viral transduction, cell lines were first grown to 70-80 % confluency in a six-well plate format, clarified by aspiration or gentle centrifugation (300 x g, 5 min) to remove spent media, and treated with purified virus resuspended in 800 µL DMEM containing 8 µg/mL polybrene. The cells were spinfected by centrifuging the six-well plates at 1048 x g for 90 min at 30 °C before returning to the 37 °C CO_2_ incubator for another 48 h. The AQP1-miniIAA7 and AQP1-AID expressing cell lines were transduced a second time with lentivirus encoding AtAFB2 (Addgene 129718) and OsTIR1 (Addgene 72834) auxin receptor F-box proteins, which are required to bind the auxin degron tag and form a complex with ubiquitin E3 ligase for degradation. To enable sorting of doubly-transduced cells, the above genes were co-expressed with an mCherry reporter. Following lentiviral transduction, stably transduced cells were enriched by fluorescence activated cell sorting (FACS) using a Sony SH800 cell sorter and stored as cryo-stocks until further use.

### Cell viability assay

To assess potential toxicity resulting from shield-1 treatment, cells were assayed for MTT reduction (CellTiter 96® Non-radioactive Cell Proliferation Assay) and intracellular ATP content (CellTiter-Glo®). For each assay, cells were grown to 80-90 % confluency in 96-well plates and incubated with the assay reagents as per the manufacturer’s protocol. Absorbance (MTT reduction) and luminescence readings (ATP content) were obtained using a plate reader (Tecan Spark). Integration time for luminescence measurements was set to 1 s.

### Western blotting

In preparation for western blotting, membrane fractions were isolated from cells using ProteoExtract® native membrane protein extraction kit and concentrated using 10 kDa Amicon® Ultra-15 Centrifugal Filters. For immunoblotting of cytoplasmic extracts, cells were directly lysed using RIPA buffer. Total protein concentration in the membrane fraction and cell lysate was estimated using BCA assay to ensure that similar amounts of total sample were probed by western blotting. Next, 20 µL of cell lysate was denatured by mixing with an equal volume of 2X Laemmli buffer supplemented with 5 % beta-mercaptoethanol and incubated at room temperature for 16 h. For the membrane fraction, denaturation was performed by boiling at 95 °C for 8 minutes. The denatured extract was resolved by electrophoresis in an SDS-PAGE gel at 90 V for 90 min at 4 °C and transferred to a polyvinylidene difluoride membrane using a Biorad Trans-Blot® Turbo™ transfer system. The membrane was blocked for 1 h in 50 mL of Tris-Buffered Saline containing 0.1 % (w/v) Tween® 20 (TBS-T) and supplemented with 5 % (w/v) non-fat dried milk. Subsequently, the membrane was incubated overnight at 4°C in 50 mL of TBS-T containing 1 µg/mL mouse anti-FLAG primary antibody and 5% (w/v) non-fat dried milk. The membrane was washed three times using 50 mL of TBS-T with an incubation time of 15 min per wash. The membrane was then treated with 50 mL TBS-T containing 0.5 µg/mL horseradish peroxidase-conjugated goat anti-mouse IgG secondary antibodies and 5% (w/v) non-fat dried and further washed with 50 mL of TBS-T three times as before. Finally, the membrane was incubated with Clarity™ Western ECL Substrate and imaged using an iBright™ FL1500 imaging system (Thermo Fisher Scientific).

### Immunofluorescence imaging

LSAqp1-transduced CHO cells were grown in 35 mm #1.5 glass bottom dishes (Ibidi, Cat. No. 81218-200) to a confluency of 70-80%. After aspirating the medium, cells were fixed by incubating in 1 mL 4% paraformaldehyde for 15 minutes at room temperature inside a fume hood. Cells were rinsed three using 2 mL sterile phosphate buffered saline (PBS) with an incubation time of 5 minutes per wash. The cells were then blocked by incubation with blocking solution (2% BSA, 5% goat serum in PBS) for 1 hour. Subsequently, the cells were incubated for 18 hours at 4°C in a rotary shaker (set at 200 r.p.m) with 1 mL blocking solution containing 5 µg/mL mouse anti-FLAG primary antibody. primary antibody solution. The cells were washed three times using 2 mL PBS with an incubation time of 5 minutes for each wash. Finally, the cells were incubated with 1mL of PBS containing Alexa Fluor 647 conjugated goat anti-mouse IgG secondary antibody (2 µg/mL) and 2% bovine serum albumin (BSA). The plate was foil wrapped, incubated in a rotary shaker (200 r.p.m) at room temperature for 2 hours, and washed two times using 2 mL PBS with an incubation time of 5 minutes at each wash step. Next, the cells were incubated for 5 minutes at room temperature with DAPI (4′,6-diamidino-2-phenylindole) lactate salt (2.5 µg/mL in PBS) to stain the nucleus, washed twice with 1 mL PBS (3 minutes incubation per wash step), and finally overlaid with 1 mL PBS in preparation for confocal imaging.

Confocal microscopy was performed using a Leica Dmi8 SP8 resonant scanning microscope equipped with an HC PL APO 63x/1.40 oil-immersion objective using Type F immersion oil. A 405 nm laser line was used for DAPI excitation and the emitted light was detected between 410 nm and 483 nm. A 653 nm laser line was used for Alexa Fluor 647 excitation and the emitted light was detected between 665 nm and 779 nm. Leica Application Suite X (LAS X) was used for image acquisition and Fiji was used for image analysis.

### In vitro MRI and differential imaging

In preparation for MRI, cells were seeded such that they reached confluency in approximately 48 h. 24 h post-seeding, cells were treated with a small-molecule ligand matched to the degron-type expressed by the cell: 10 µM trimethoprim (degron derived from dihydrofolate reductase), 1 µM 4-hydroxytamoxifen (degron derived from estrogen receptor), 1 µM shield-1 (FKBP12^F36V/L106P^ degron), 500 µM indole-3-acetic acid (mini-IAA7 and AID), and 3 µM asunaprevir (SMASh). For the proteasomal and lysosomal inhibition experiments, cells were treated 24 hours post seeding with 100 µM chloroquine or 10 µM MG132. Control cohorts were prepared by treating cells with the respective vehicle solution, i.e. ethanol (for indole-3-acetic acid) and 0.1 % dimethyl sulfoxide (for all the other compounds). Adherent cells were harvested by trypsinization, centrifuged at 350 x g, resuspended in 200 µL sterile PBS and transferred to 0.2 mL tubes. For the mixed cell experiments, the two cell-types were cultured separately, counted using a hemocytometer, mixed by gentle pipetting, centrifuged as before, and transferred to 0.2 mL tubes. The tubes were centrifuged at 500 x g for 5 min to form a cell pellet and placed in water-filled agarose (1 % w/v) molds housed in a 3D-printed MRI phantom. Imaging was performed using a 66 mm diameter coil in a Bruker 7T vertical-bore MRI scanner. Stimulated echo diffusion-weighted images of cell pellets were acquired in the axial plane using the following parameters: echo time, T_E_ = 18 ms, repetition time, T_R_ = 1000 ms, gradient duration, δ = 5 ms, gradient separation, Δ = 300 ms, matrix size = 128 × 128, field of view (FOV) = 5.08 × 5.08 cm^2^, slice thickness = 1-2 mm, number of averages = 5, and typically 4 b-values in the range of 1000-3000 s/mm^2^. Diffusion-weighted intensity at a given b-value was estimated by computing the average intensity of all voxels inside a region of interest (ROI) encompassing the axial cross section of a cell pellet. The slope of the logarithmic decay in mean signal intensity as a function of b-value was used to calculate the apparent diffusivity (D). In some cases, apparent diffusivity was computed for each voxel in an ROI and used to generate a diffusion map. Least-squares regression fitting was performed using the fitnlm function in Matlab (R2022b).

Differential imaging was performed by voxel-wise subtraction of diffusion-weighted images or diffusion maps of cells belonging to various treatment groups (e.g. shield-1, shield-1 + doxycycline, shield-1 + trimethoprim) from corresponding images of vehicle-treated cells. The difference images were denoised using a median filter and pseudo colored according to an 8-bit color scale. Image subtraction was performed in Matlab (R2022b), while median filtering and pseudo coloring were implemented in Fiji.

### Kinetic studies

For the kinetic studies, pellets of LSAqp1-transduced CHO cells were collected at multiple time points (t = 0, 6, 12, and 24 h) following treatment with shield-1. In vitro MRI (i.e. imaging of cell pellets) is an end-point technique. Therefore, out of practical necessity, cells corresponding to each time point were harvested from different plates, which were all seeded in a similar manner from cryo-stocks. Diffusion-weighted imaging and whole-cell diffusivity measurements were performed exactly as described above.

### Mouse tumor xenograft

CHO cells expressing LSAqp1 or Aqp1-only (viz. without a degron) were grown to 80-90 % confluency, harvested by trypsinization, centrifuged at 300 x g for 5 min, and resuspended in 100 µL sterile PBS. The resuspended cells were mixed with an equal volume of Matrigel®, loaded into 1 mL sterile syringes fitted with 25G needles, and subcutaneously injected in the left and right hind limbs of 5-7 weeks old female NOD/SCID/γ-mice (Jackson Labs Strain No. 005557). Tumors were measured daily using calipers and tumor volume was calculated as 0.52 x (short axis)^2^ x (long axis)^2^. All protocols were approved by the Institutional Animal Care and Use Committee of the University of California, Santa Barbara.

### In vivo MRI

Once the subcutaneous tumors reached a volume of 100-300 mm^3^ (17-21 days post-implantation), mice were imaged using a Bruker 7T vertical-bore MRI scanner. Mice were first anesthetized using 2-3 % isoflurane and secured in an animal cradle with medical tape to ensure proper positioning of tumors and to minimize motion-induced artifacts. The cradle was inserted in the MRI scanner concentrically with a 40 mm diameter imaging coil. Respiration rate and body temperature were monitored throughout the imaging session using a pneumatic cushion (Biopac, Goleta, CA, USA) and a fiber optic rectal probe (Opsens) respectively. Body temperature was maintained at 37 °C using warm air flow with feedback temperature control. To induce Aqp1 and LSAqp1 expression in the tumor cells, mice were intraperitoneally injected with 50 mg/kg doxycycline 24 h before imaging.

The location of the tumors was first determined using a high-resolution multi-slice axial scan with the following parameters: echo time, T_E_ = 14.22 ms, repetition time, T_R_ = 660.2 ms, RARE factor = 4, matrix size = 256 × 256, field of view (FOV) = 3.5 × 3.5 cm^2^, slice thickness = 1 mm, number of averages = 10, and number of slices = 10. Diffusion-weighted images were subsequently obtained using a stimulated echo pulse sequence with the following parameters: echo time, T_E_ = 18 ms, repetition time, T_R_ = 1000 ms, gradient duration, *δ* = 5 ms, gradient interval, Δ = 300 ms, matrix size = 96 × 96, field of view (FOV) = 3.5 × 3.5 cm^2^, number of averages = 18, and effective b-values of 1255 and 3377 s mm^-2^. Next, mice were intraperitoneally injected with 10 mg/kg shield-1 dissolved in a mixture of 50% *N,N*-dimethylformamide and 50% 9:1 PEG-400: Tween®-80.

Approximately 6 h after injecting shield-1, the mouse was returned to the imaging cradle. The post-injection scan was acquired using the same procedure as before, taking care to position the axial slice to align as closely as possible with the pre-injection tumor plane. While manual alignment of pre- and post-injection slices is considerably challenging, we note that this step can be greatly simplified through the use of commercially available automated mouse positioning systems. To calculate apparent diffusivity, ROI(s) were manually drawn (using Fiji) in the tumor volume. Mean intensities were calculated for each ROI and diffusivity was estimated from the decay in signal intensity versus b-value.

Differential images were generated by performing rigid-body registration of pre- and post-injection images of the tumor ROIs followed by voxel-wise subtraction of signals. Rigid-body registration was implemented using the imregister function in Matlab (R2022b). The ensuing difference image was pseudo colored and overlaid on a high-resolution anatomical scan obtained in the same plane as the diffusion-weighted image.

### Statistical analysis

Data are summarized by their mean and standard deviation obtained from multiple (*n* ≥ 3) biological replicates and compared using a Student’s t-test. Quality of model-fitting (for estimating diffusivity) was judged based on the regression coefficient and inspection of the 95 % confidence interval and residuals. All tests are 2-sided and a *P* value of less than 0.05 taken to indicate statistical significance.

## Author Contributions

AM conceived the study. AM and JY designed experiments. JY and YH performed all experiments with help from LB, EL, HL, ANC, JW, and ADM. JY, YH, and AM performed all data analysis. AM wrote the manuscript with inputs from all authors. AM supervised the research.

## Acknowledgements

We thank members of the Mukherjee lab for helpful discussions. Dr. Jerry Hu (UC, Santa Barbara) is gratefully acknowledged for assistance with setting up the MRI workflow. This research was supported by the National Institutes of Health (R35-GM133530, R21-EB033989, R03-DA050971, and R01-NS128278 to A.M.), the U.S. Army Research Office via the Institute for Collaborative Biotechnologies cooperative agreement W911NF-19-D-0001-0009 (A.M.), and a NARSAD Young Investigator Award from the Brain & Behavior Research Foundation (A.M.). This project has been made possible in part by a grant from the Chan Zuckerberg Initiative DAF, an advised fund of Silicon Valley Community Foundation. J.Y. gratefully acknowledges support from the Chair’s Fellowship from UC Santa Barbara (Department of Chemistry). L.B. acknowledges support from the National Science Foundation Graduate Research Fellowships Program. All MRI experiments were performed at the Materials Research Laboratory (MRL) at UC, Santa Barbara. The MRL Shared Experimental Facilities are supported by the MRSEC Program of the NSF under Award No. DMR 1720256; a member of the NSF-funded Materials Research Facilities Network. We acknowledge the use of the Neuroscience Research Institute-MCDB Microscopy Facility and the Resonant Scanning Confocal supported by NSF MRI grant 1625770.

