## Supplementary Information for "Engineering ligand stabilized aquaporin reporters for magnetic resonance imaging"

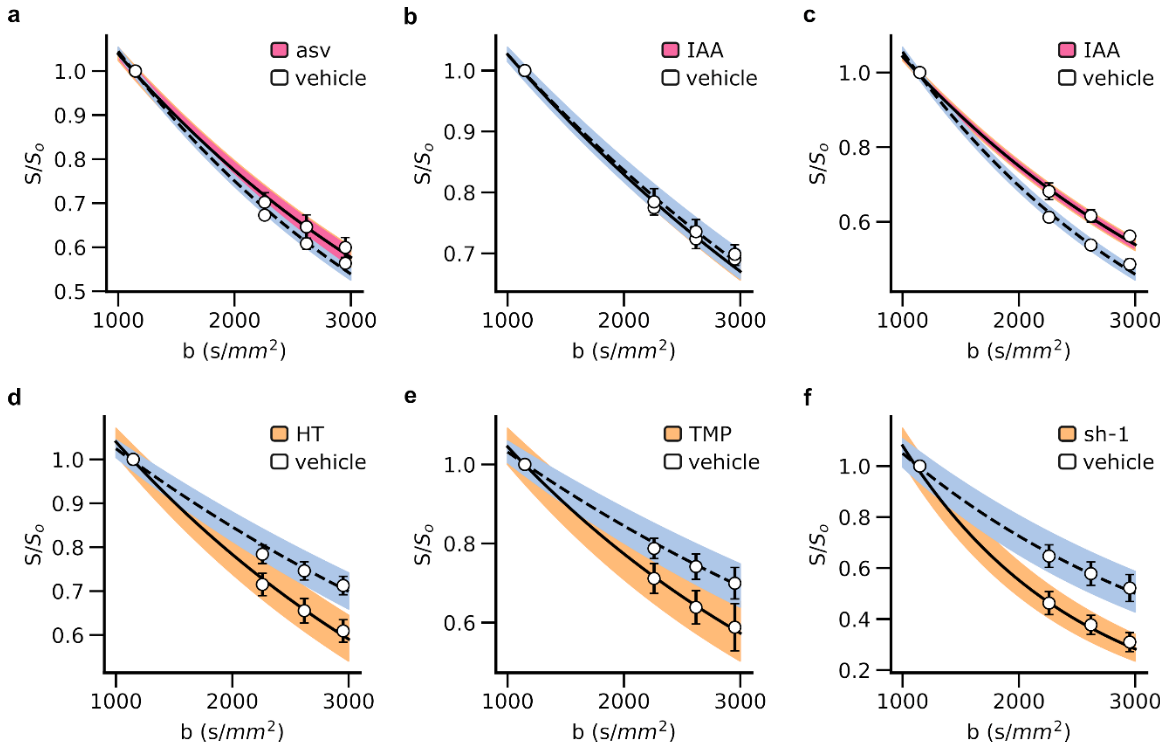


**Supplementary figure 1: Diffusivity measurements in CHO cells engineered to express various degron-tagged Aqp1 reporters.** Ligand-induced degradation (LID) constructs comprised Aqp1 fused to the following degrons: **a)** small molecule-assisted shutoff (SMASh), which responds to 3 µM asunaprevir (asv)[1]; **b)** auxin-inducible degron (AID), which responds to 0.5 mM indole-3-acetic acid (IAA)[2]; and **c)** a truncated version of AID (amino acids 37-104), known as mini-IAA7[3], which is also responsive to IAA (0.5 mM). Ligand-induced stabilization (LIS) constructs comprised Aqp1 fused to the following degrons: **d)** a degron derived from the human estrogen receptor[4], which is stabilized in response to 1 µM 4-hydroxy tamoxifen (HT); **e)** a degron derived from dihydrofolate reductase[5], which responds to 10 µM trimethoprim (TMP); and **f)** FKBP12$^{F36V/L106P}$ degron, which is stabilized by 1 µM shield-1[6,7]. Error bars represent the standard deviation ($n \geq 4$ biological replicates). Solid and dotted lines represent the best-fit plots of the decay in signal intensity (S) with diffusion weighting (effective b-value) for cells treated with the cognate ligand and mock-treated cells, respectively. The shaded region represents the 95 % confidence band estimated from the replicate measurements of the diffusion coefficients.

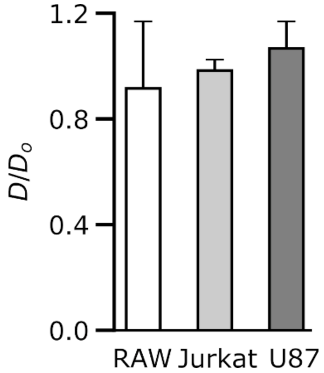


**Supplementary figure 2: Specificity of shield-1 modulation.** Treatment with shield-1 (1 μM) did not alter diffusivities in cells engineered to express Aqp1 without the $FKBP12^{F36V/L106P}$ degron ($P$ > 0.2, 2-sided t-test). $D/D_o$ represents the fold-change in the diffusion coefficients of cells treated with shield-1 relative to those of cells treated with vehicle (0.1 % DMSO). Error-bars represent the standard deviation ($n$ = 4 biological replicates).

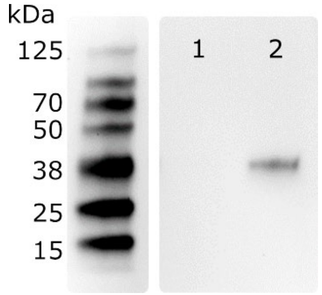


**Supplementary figure 3**. **Western blotting of membrane fractions from LSAqp1-transduced U87 cells**. Membrane fractions were prepared from U87 cells stably transduced with LSAqp1 incorporating a FLAG epitope at the N-terminus. LSAqp1 expression was probed by immunostaining using an anti-FLAG antibody. Lane 1: vehicle-treated (0.1 % DMSO) cells; lane 2: cells treated with 1 μM shield-1.

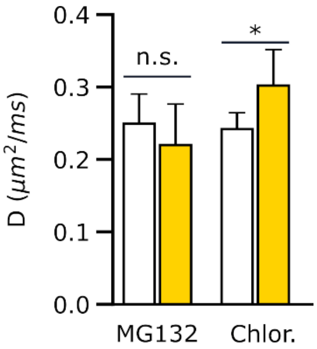


**Supplementary figure 4. Putative mechanism of LSAqp1 degradation**. Overnight incubation of LSAqp1-transduced CHO cells with 10 μM MG132 (proteasomal inhibitor) did not induce a significant change in diffusion rates. In contrast, the incubation of cells with 100 μM chloroquine (a lysosomal inhibitor) generated a modest but statistically significant increase in the diffusion coefficient of LSAqp1 cells. Error bars represent the standard deviation. * denotes $P < 0.05$ and n.s. denotes non-significant ($P \geq 0.05$). $P$-values were computed using a 2-sided Student's t-test (unpaired).

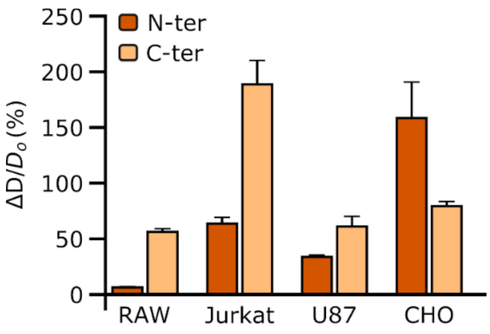


**Supplementary figure 5. Shield-1 modulation of Aqp1 with an N-terminal FKBP12$^{F36V/L106P}$ degron**. Compared to LSAqp1, incubation with shield-1 (1 µM) elicited a smaller change in the diffusion coefficient in RAW264.7, U87, and Jurkat cells transduced to express the N-terminal FKBP12$^{F36V/L106P}$-Aqp1 construct but a larger response in identically transduced CHO cells. Error bars represent the standard deviation ($n \geq 3$ biological replicates).

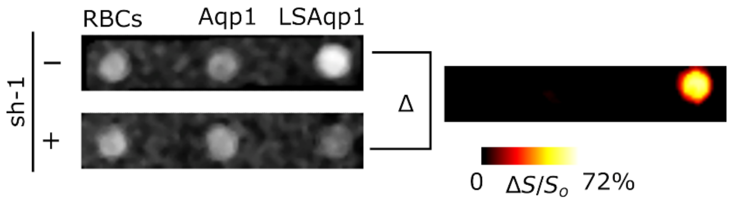


**Supplementary figure 6. Background-free imaging based on small-molecule modulation of LSAqp1**. Voxel-wise subtraction of diffusion-weighted images acquired with shield-1 from those obtained without shield-1 permits LSAqp1 expression to be disambiguated from other sources of rapid diffusion such as red blood cells, Aqp1-expressing cells, and tissue background-mimicking agarose. The differential image was denoised using a median filter and displayed as a pseudo-colored “hotspot”.

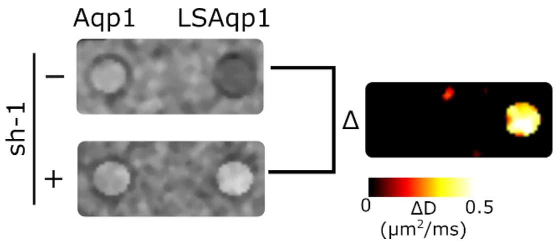


**Supplementary figure 7**. **Background-free imaging based on diffusion maps**. In a diffusion map, each voxel is represented by its diffusion coefficient, which is computed from the decay of the diffusion-weighted signal intensity with b-value. Voxel-wise subtraction of diffusion maps acquired in the presence of shield-1 from diffusion maps of vehicle-treated cells revealed signals specific to the LSAqp1-expressing cell population.  The difference image is denoised using a median filter and displayed as a pseudo-colored “hotspot”.

**a**

| sh-1 | $T_x$ | Output |
|---|---|---|
| 0 | 0 | 0 |
| 1 | 0 | 0 |
| 0 | 1 | 0 |
| 1 | 1 | 1 |

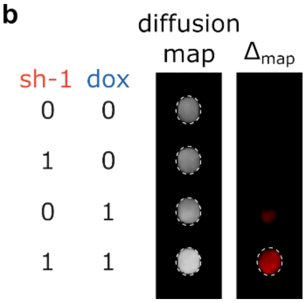


**Supplementary figure 8. Chemically gated imaging of transcriptional activity using LSAqp1**. **a)** Truth table for AND gate design for chemically gated imaging of promoter activity using LSAqp1. $T_X$ represents the transcriptional activity induced by doxycycline. **b)** Diffusion maps are generated by representing each voxel with its diffusivity. Differential images were generated by voxel-wise subtraction of diffusion maps acquired in the presence of shield-1 and/or doxycycline from diffusion maps of untreated cells. A positive signal was observed only when promoter activity was accompanied by shield-1.

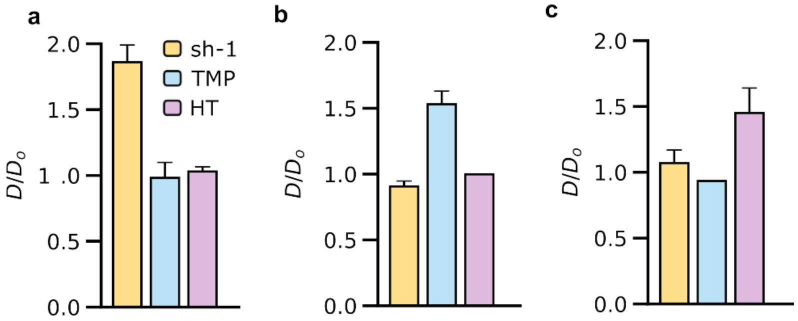


**Supplementary figure 9: Ligand-stabilized Aqp1 reporters are mutually orthogonal.** LSAqp1, Aq-dhfr, and Aq-ER exhibit non-overlapping ligand requirements. **a)** Treatment with shield-1, but not trimethoprim (TMP) or 4-hydroxytamoxifen (HT), increases the diffusion coefficient of LSAqp1-transduced cells. **b)** Treatment with TMP, but not shield-1 or HT, increases the diffusivity of Aq-dhfr-transduced cells. **c)** Treatment with HT, but not shield-1 or TMP, increases the diffusion coefficient of Aq-ER-transduced cells. All fold-changes were measured relative to reporter-transduced cells treated under identical conditions with 0.1 % DMSO.

**Table S1. Plasmids engineered and used in this study.**

| Plasmid | Main constructs encoded | Notes |
|---|---|---|
| pJY22 | Flag-Aqp1-IRES-EGFP | Constitutive expression of degron-free Aqp1 from EF1α promoter |
| pJY02 | Flag-Aqp1-AID-IRES-EGFP | Expresses Aqp1 tagged to the auxin-inducible degron (AID) |
| pJY09 | OsTIR1-IRES-mCherry | Expresses OsTIR1, an auxin receptor F-box protein that is needed to activate AID |
| pJY16 | Flag-Aqp1-SMASh-IRES-EGFP | Expresses Aqp1 tagged to the small-molecule assisted shutoff degron (SMASh) |
| pJY19 | Flag-Aqp1-IAA7-IRES-EGFP | Expresses Aqp1 tagged to miniIAA7 |
| pJY18 | AtFB2-IRES-mCherry | Expresses AtFB2, an auxin receptor F-box protein needed to activate miniIAA7 |
| pJY20 | Flag-Aqp1-DHFR-IRES-EGFP | Constitutive expression of Aqp1 tagged to the dihydrofolate reductase-based degron (DHFR) |
| pJY21 | Flag-Aqp1-ER-IRES-EGFP | Constitutive expression of Aqp1 tagged to the human estrogen receptor-based degron (ER) |
| pJY23 | Flag-Aqp1-FKBP12-IRES-EGFP | Constitutive expression of Aqp1 tagged to the $FKBP12^{F36V/L106P}$ degron (FKBP12) |
| pJY25 | Flag-FKBP12-Aqp1 -IRES-EGFP | Constitutive expression of Aqp1 tagged to the $FKBP12^{F36V/L106P}$ degron at its N-terminus |
| pJY12 | Flag-Aqp1-FKBP12-IRES-EGFP | Expresses Aqp1 tagged to the $FKBP12^{F36V/L106P}$ degron (C-terminus) from a doxycycline-inducible promoter |
| pJY00 | Flag-Aqp1-IRES-EGFP | Expresses degron-free Aqp1 from a doxycycline-inducible promoter |
| pJY23_flag7 | Aqp1(flag)-FKBP12-IRES-EGFP | Constitutive expression of Aqp1 with a FLAG tag inserted in an extracellular loop (between Q43 and T44) |
| pPackaging | | Expresses proteins for lentiviral packaging |
| pVSV-G | | Expresses the VSV-G protein for broad lentiviral tropism |
